# High-order interdependencies in the aging brain

**DOI:** 10.1101/2020.03.17.995886

**Authors:** Marilyn Gatica, Rodrigo Cofré, Pedro A.M. Mediano, Fernando E. Rosas, Patricio Orio, Ibai Diez, S.P. Swinnen, Jesus M. Cortes

## Abstract

Brain interdependencies can be studied either from a structural/anatomical perspective (“structural connectivity”, SC) or by considering statistical interdependencies (“functional connectivity”, FC). Interestingly, while SC is typically pairwise (white-matter fibers start in a certain region and arrive at another), FC is not; however, most FC analyses focus only on pairwise statistics and neglect high-order interactions. A promising tool to study high-order interdependencies is the recently proposed *O-Information*, which can quantify the intrinsic statistical synergy and redundancy in groups of three or more interacting variables. In this paper we used the O-Information to investigate how high-order statistical interdependencies are affected by age. For this, we analised functional magnetic resonance imaging (fMRI) data at rest obtained from 164 healthy participants, ranging from 10 to 80 years old. Our results show that older subjects (age ranging from 60 to 80 years) exhibit a higher predominance of redundant dependencies than younger subjects; moreover, this effect seems to be pervasive, taking place at all interaction orders. Additionally, we found that these effects are highly heterogeneous across brain regions, and suggest the existence of a “redundancy core” formed by the prefrontal and motor cortices – thus involving functions such as working memory, executive and motor functions. Our methodology to assess high-order interdependencies in fMRI data has unlimited applications. The code to calculate these metrics is freely available.

## 1. Introduction

Deepening our understanding of the anatomical and functional correlates of aging in the brain has great scientific, medical, and social implications, particularly as projections suggest that the world’s elderly population might nearly double from 2015 to 2050 [1]. Aging causes a systemic decline at multiple scales, from biological to cognitive and psychosocial levels. Aging tends to disrupt the circadian behavior and sleep cycles, which in turn causes a decrease in sleep quality [2]. It also affects multiple cognitive domains such as information processing speed, working memory, executive functions, and reasoning [3], as well as several mental health conditions (such as anxiety and depression) [4].

This age-related cognitive decline occurs in parallel with well-established variations of brain morphology. Its total volume is known to increase from childhood to adolescence in about 25% on average, remains constant for the three following decades and finally decays back to childhood size at late ages [5]. Strikingly, the amount of atrophy is not homogeneous across brain areas, but some anatomical regions are more affected than others; well-known aging-targeted structures are the hippocampus [6] and the prefrontal cortex [7].

The development of resting state functional magnetic resonance imaging (fMRI) has shown that age affects the functional connectivity (FC) of large-scale brain networks, specifically between anterior and posterior regions, including superior and middle frontal gyrus, posterior cingulate, middle temporal gyrus, and the superior parietal region [8, 9]. Other studies have shown that age can also increase FC between several regions, possibly indicating compensation or pathological activation [10, 11, 12, 13].

An important limitation of the above FC studies is that their analyses are restricted to pairwise functional connectivity, ignoring possible high-order effects. High-order interactions allow us to distinguish redundancy and synergy dominated interactions, which have shown to play key roles in neural dynamics [14, 15, 16, 17, 18, 19]. A first study on high-order interactions and aging was reported in Ref. [20], which showed significant changes in synergies and redundancies in triple interactions along the lifespan, and a redundant role of the default mode network. The effect of aging on interactions beyond triple relationships is, to the best of our knowledge, still an unexplored territory.

In this paper, we build upon Ref. [20] and study the effects of aging on the high-order interactions in the human brain, with a special focus on interdependencies between four or more brain regions. To this end, we employ the recently proposed *O-Information* [21], which can be considered to be revision of the measure of neural complexity proposed by Tononi, Sporns and Edelman [22] under the light of Partial Information Decomposition [23]. In particular, the O-Information captures the balance between redundancies and synergies in arbitrary sets of variables, thus extending the properties of the interaction information of three variables to larger sets [24]. Redundancy is here understood as an extension of the conventional notion of correlation to more than two variables, in which each variable has a “copy” of some common information shared with other variables [25]; an example of extreme redundancy is full synchronisation, where the state of one signal allows one to predict the state of any other. In contrast, synergy corresponds to statistical relationships that regulate the whole but not the parts [21, 26, 27]. Synergies allow the coexistence of local independency and global cohesion, being conjectured to be instrumental for high order brain functions, while redundancy – including cases of high synchronization such as deep sleep or epileptic seizures – would make brains less well-suited for them [22, 28].

In order to investigate how the high-order informational organization of the brain changes with age, we studied synergistic and redundant interactions for different interaction orders from data of functional magnetic resonance imaging (fMRI) in resting state from a cohort of 164 healthy volunteers ranging from 10 to 80 years old. Our results show that the interdependencies seen in older subjects are more redundancy-dominated than those in younger participants, at all interaction orders. When studying how these effects are distributed topographically, we found a “redundancy core” composed by brain regions that participate in the most redundancy-dominated arrangements for all interaction orders.

## 2. Materials and methods

### 2.1. Participants

A sample of *N* = 164 healthy volunteers participated in the study with an age ranging from 10 to 80 years (mean age 44.35 years, SD 22.14 years). Informed consent was obtained before testing. The study was approved by the local ethics committee for clinical research and was performed in accordance with the Declaration of Helsinki.

We grouped the participants into four age groups *I*_*i*_ for *i* ϵ {1, …, 4}. More specifically, *I*_1_ consists of 30 subjects with ages ranging from 10-20 years, *I*_2_ of 46 subjects from 20-40 years, *I*_3_ of 29 subjects from 40-60 years and *I*_4_ of 59 subjects from 60-80 years.

### 2.2. Experimental

Image acquisition per participant was performed in an MRI Siemens 3T MAGNETOM Trio MRI scanner with a 12-channel matrix head coil. The anatomical data was acquired as a high-resolution T1 image with a 3D magnetization prepared rapid acquisition gradient echo: repetition time (RT) = 2300 ms, echo time (ET) = 2.98 ms, voxel size =1×1×1.1 mm^3^, slice thickness = 1.1 mm, field of view =256×240 mm^2^, 160 contiguous sagittal slices covering the entire brain and brainstem. Resting state functional data was acquired with a gradient echo-planar imaging sequence over a 10 min session using the following parameters: 200 whole-brain volumes with TR/TE = 3000/30 ms, flip angle =90, inter-slice gap =0.28 mm, voxel size =2.5×3×2.5 mm^3^, 80×80 matrix, slice thickness =2.8 mm, 50 oblique axial slices, interleaved in descending order.

Image preprocessing was performed following a similar procedure to that in Ref. [20]. After preprocessing, each subject was represented by a set of 2514 time series of fMRI signal, each corresponding to a region of interest (ROI), here built as the average signal within a super-voxel consisting of about 35 voxels of size 2.5×3×2.5 mm^3^ [29].

### 2.3. Brain Hierarchical Atlas

The Brain Hierarchical Atlas (BHA) is a brain parcellation constructed by a hierarchical agglomerative clustering combining both functional and structural data [29]. In this study, the original 2514 time series per subject are grouped in 20 modules or brain regions, which is the one that maximizes the matching between brain functional modules and structural ones [29]. As the fMRI signals of the ROIs belonging to the same module are similar, we used the average of the ROIs belonging to the same module as the representative signal of that module. Therefore, the brain dynamics for each subject was reduced into 20 time series (see Fig. 1).

**Figure 1:**
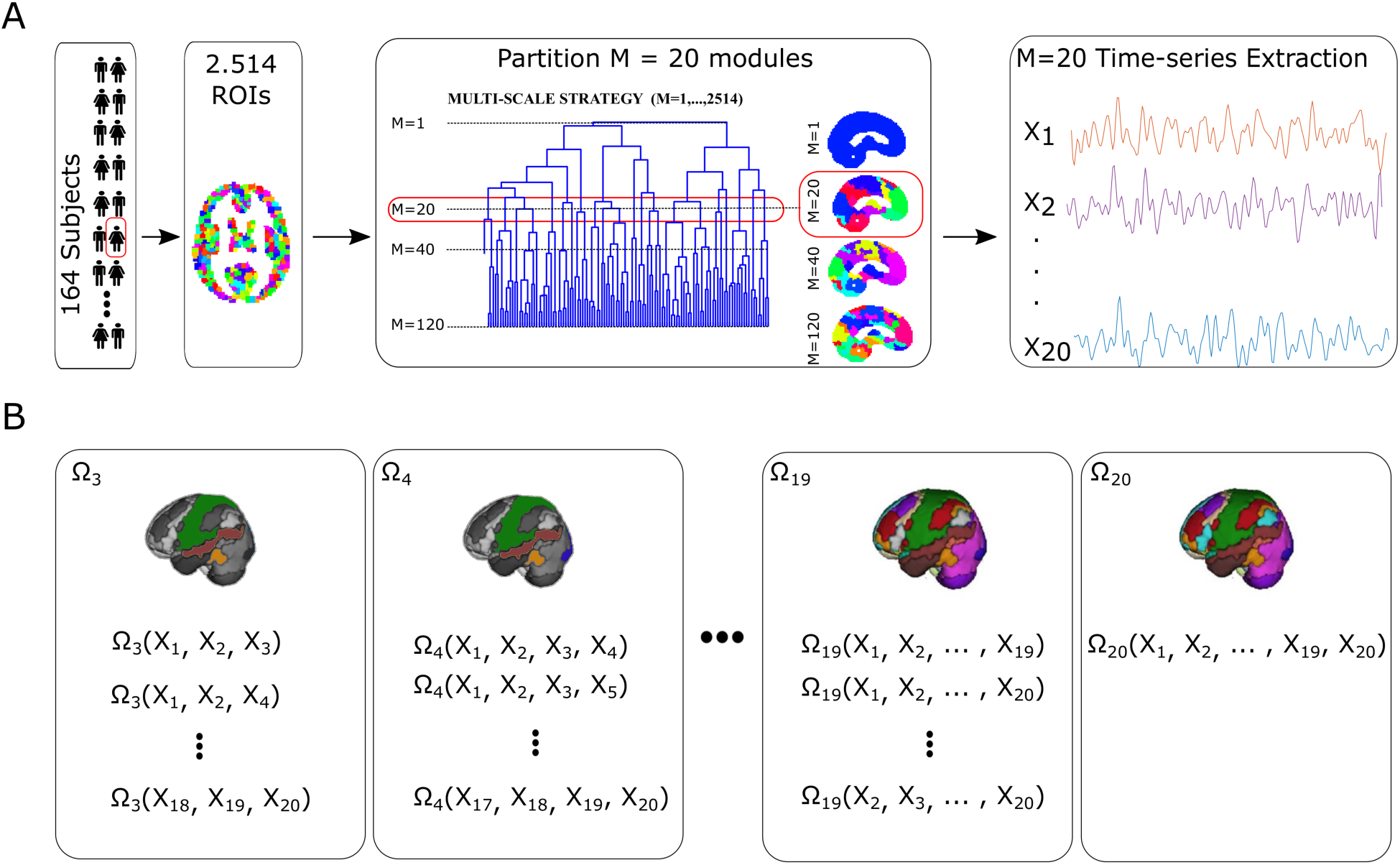
Method overview. **A:** 164 subjects were included in this study. For each of them, 2514 fMRI signals were obtained, and parcellated into 20 regions using the Brain Hierarchical Atlas (BHA). All ROIs corresponding to each module were averaged to obtain one representative signal per module for each subject. **B:** For each subject and interaction order we computed for each n-plet their O-Information and S-Information. These values were used for further analysis, as described in detail in the Appendix.

### 2.4. Multivariate information-theoretic metrics

The O-Information [21] (shorthand for *information about organisational structure*) is an attempt to operationalise the original desiderata proposed in Ref. [22] while overcoming some of the shortcomings of their original proposal (see e.g. [30, 31]). Denoted by Ω(***X***^*n*^), the O-Information is a real-valued measure whose sign serves to discriminate between redundant and synergistic components. In effect, Ω(***X***^*n*^) *>* 0 corresponds to redundancy-dominated interdependencies, while Ω(***X***^*n*^) *<* 0 characterises synergy-dominated ones. When applied over groups of three variables, Ω(***X***^3^) is equal to McGill’s *interaction information* [24]; however, their values differ for larger system sizes. Importantly, Ω(***X***^*n*^) (but not the interaction information) keeps the ability to discriminate between redundancy-and synergy-dominated systems for *n* > 3.

In addition to the O-Information, we also employ the S-information [21] – denoted by Σ(***X***^*n*^) – to quantify the strength of multivariate correlations. The S-information is a natural complement of the O-Information: while the former quantifies the overall strength of the interdependencies, the latter determines whether they are synergistic or redundant in nature. Furthermore, following Ref. [20], we take the positive and negative parts of the O-Information, denoted as 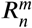 and 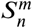, as heuristic metrics of redundancy and synergy.

The Appendix provides details about how these metrics are calculated.

### 2.5. Interaction order, redundancy and synergy

We quantified the balance between redundancy and synergy in groups of brain regions by calculating the O-Information (in nats) of their corresponding fMRI signals. More precisely, we consider the set of *M* = 20 BHA modules, and for each collection of indices ***α*** = (*α*_1_, …, *α*_*n*_) ⊆ {1, …, 20} of size *n*, we computed the quantity 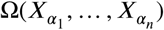 following Eq. (1) in the Appendix. We call ***α*** an *n*-plet (see Fig. 1), with *n* being its *interaction order*. Importantly, as detailed in the Appendix, the values of the O- and S-Information do not depend on the specific order of the variables within the n-plets. To compute the mean O- and S-Information at interaction order *n*, we averaged their values over all *n*-plets. Additionally, we calculate proxies for the redundancy and synergy for each module *m* and interaction order *n* using Eqs. (4) and (5) respectively, thus generalising the approach presented in Ref. [20] beyond triple interactions.

We refer to a *population value* when averaging any of the *n*-plet measures across subjects. To identify redundant brain areas, we rank all values of Ω(***X***^*n*^) in decreasing order and take the maximum value for the most redundant n-plet. To identify synergistic areas, we did the same but then take the minimal one (see Figs. 5 and 6).

### 2.6. Statistical analyses

In this study we compared the group of old participants I4 versus the combination of the three other groups (I1, I2, I3). We compared different information-based measures between different groups. Group differences were assessed by the non-parametric statistical Wilcoxon rank sum test. When appropriate, significance levels for hypothesis testing were corrected for multiple comparisons controlling the false discovery rate (FDR) following a standard Benjamini-Hochberg procedure [32].

## 3. Results

We first measured the S-Information and the O-Information per age group and interaction order. Fig. 2 shows that both the S-Information and the O-Information exhibit significant differences between the old group and the younger ones after correcting with FDR for multiple comparisons. The increase of S-information with age implies an increase of interdependencies between the various brain regions. Interestingly, the increase shown in the older population is significant at all orders, suggesting a widespread effect. The increase seen in the O-information suggest that the correlations seen in the older population are in general redundancy-dominated, becoming stronger in large orders.

**Figure 2:**
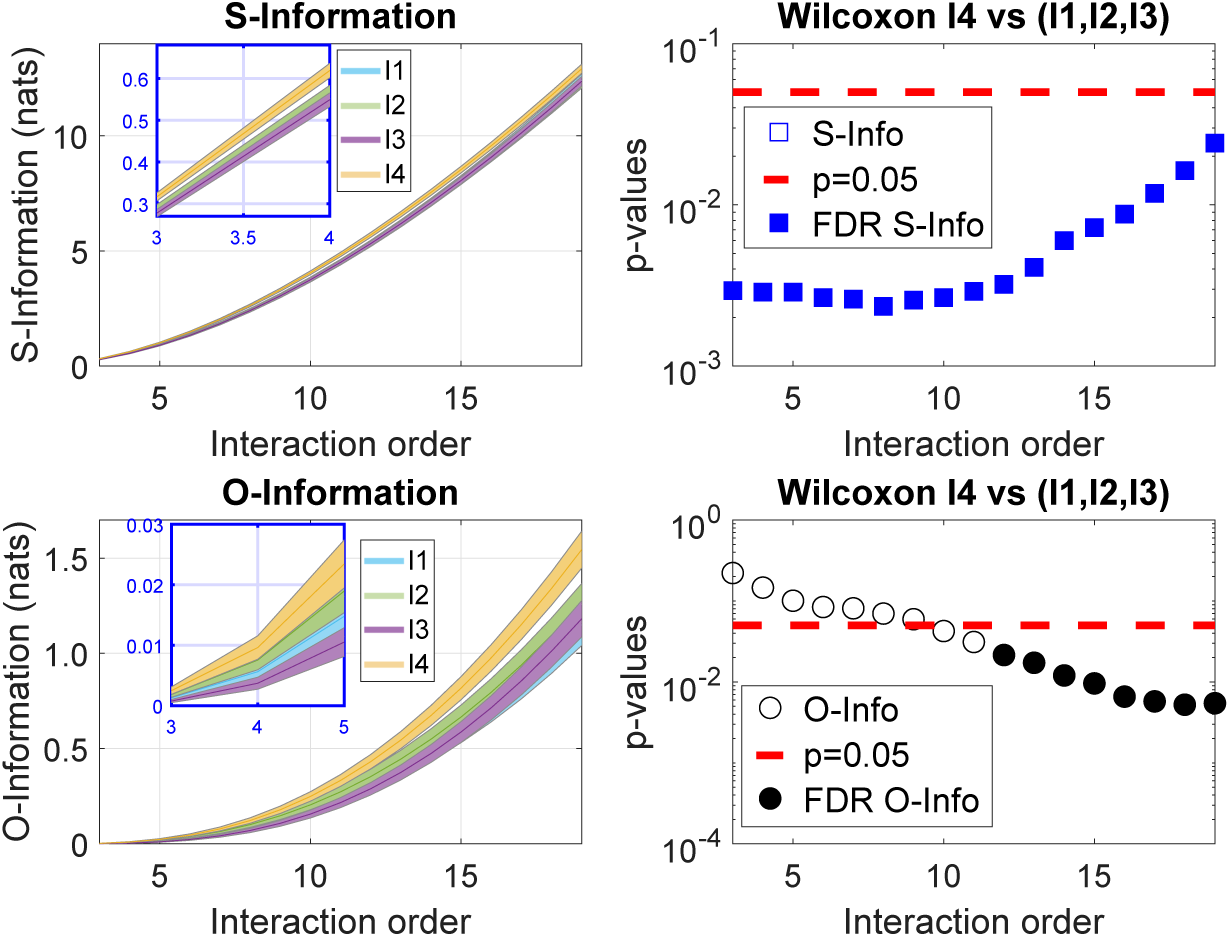
Strong high-order interdependencies in the aging brain. Both S-Information (top panel) and O-Information (bottom panel) can differentiate the high-order interdependencies of the old brains as compared to the younger ones. This is shown in the right column, where the significance after comparing the group of old people I4 vs the combination of the rest (I1, I2, I3) is represented, either uncorrected for a p value of 0.05 (red dashed line) or corrected by FDR (represented by blue-filled squares for S-Information and by black-filled circles for O-Information). The interaction order is shown on the x-axis for all plots.

By dissecting the O-Information, our results show different patterns of redundancy and synergy when increasing the interaction order (see Fig. 3). While the synergy exhibits an inverted-U (concave) shape, redundancy monotonically increases with the interaction order. Importantly, the redundancy values are much larger than the synergistic ones. Moreover, the redundancy of I4 shows significant differences from the group formed by (I1, I2, I3) for all interaction orders. For the case of synergy, while at some interaction orders the group I4 exhibit significant differences with the rest of the population, these differences do not survive the multiple comparison correction.

**Figure 3:**
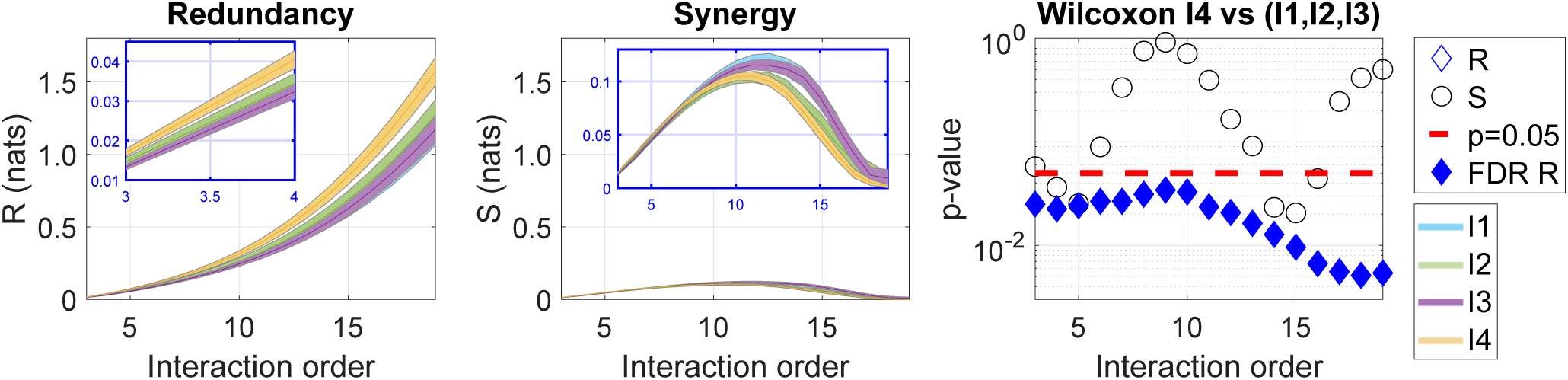
Significantly increased redundancy in older participants across all interaction orders. Average over all modules *m* = 1, …, *M* of redundancy and synergy obtained with the equations (4) and (5) for each of the groups I1, I2, I3, I4. Notice that when increasing the interaction order, and independently of age, both redundancy and synergy curves have a completely different pattern (one monotonously increasing and the other following an U-inverted curve). The right panel shows that group differences in redundancy (represented by diamonds) in older participants (I4) is significantly different from the combination of other groups (I1, I2, I3) at all interaction orders. In relation to synergy (represented by circles), none of the values survived correction for multiple comparisons at any of the interaction orders. Both diamonds and circles are filled when the value of redundancy or synergy survived correction for multiple comparisons.

When studying redundancy across brain areas, Fig. 4 shows that modules 1-3, 5, 13-15, 18-20 exhibit significant differences at all interaction orders, while the others only exhibit significant differences for large orders. This suggests the existence of a *redundancy core*, which was confirmed by later analyses. In contrast, the pattern of synergy across brain areas is highly heterogeneous, with only modules 15, 17, 18 and 19 showing significant differences.

**Figure 4:**
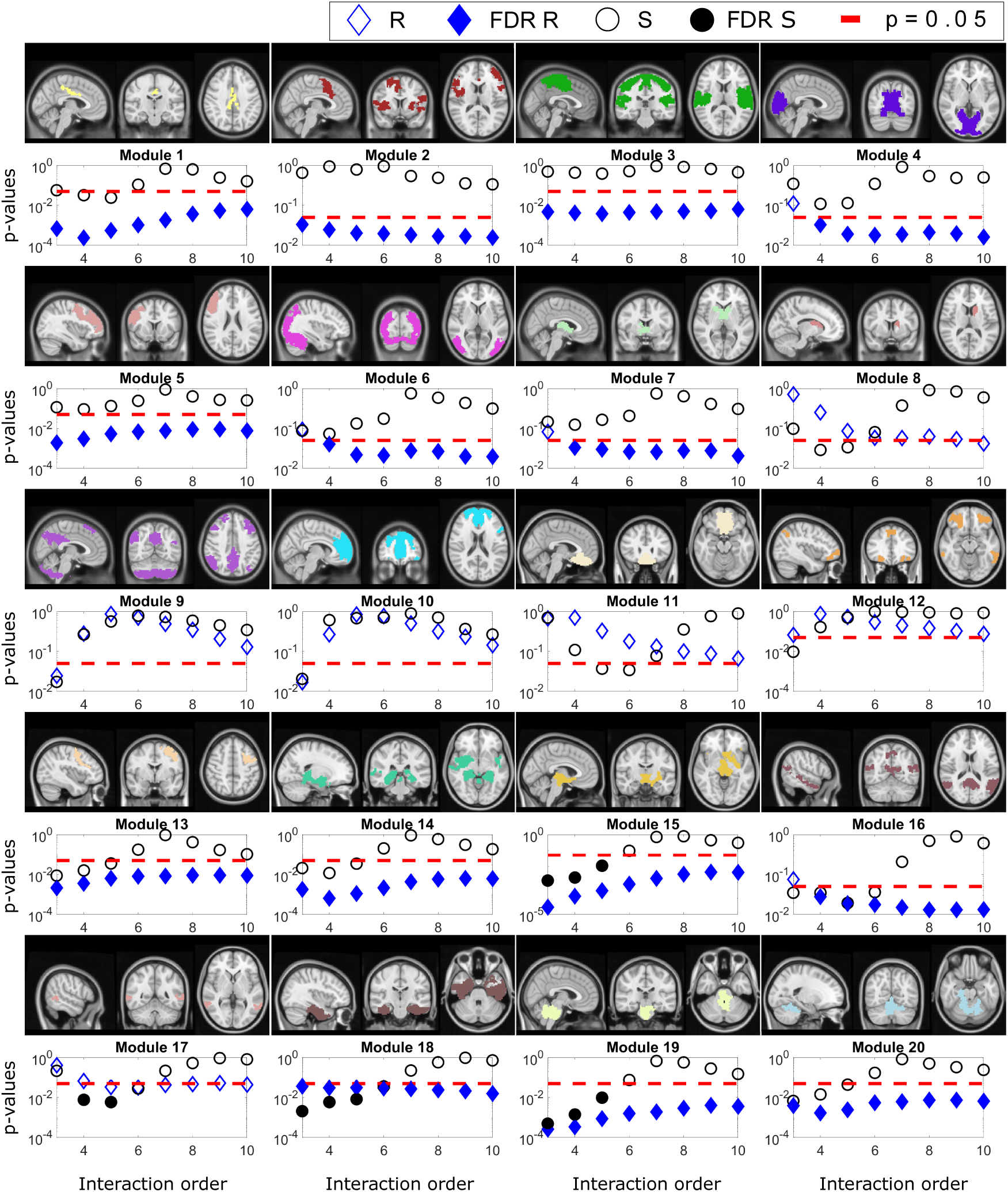
High-order redundant and synergistic interdependencies across brain areas in the aging brain. Differences between I4 and (I1, I2, I3) across brain modules, each one anatomically represented by three representative views, respectively from left to right, sagittal, coronal and axial. Similar to Fig. 3, group differences in redundancy (diamonds) and synergy (circles) are represented as a function of the interaction order. When the group differences survived correction for multiple comparisons, both diamonds and circles are filled. In relation to redundancy, there are two “classes” of modules, those where redundancy was significantly different at any order (such as modules 1-3, 5, 13-15 and 18-20) and the remaining modules for which this did not happen. Moreover, in general, it is shown that redundancy is widely different between group I4 and the rest of the population across brain areas, but not synergy, for which only modules 15, 17, 18 and 19 showed a few interaction orders with values surviving to multiple comparisons.

To confirm the existence of a redundancy core, we study the extreme values of the O-information at various interaction orders. Fig. 5 shows how modules 2, 5 and 13 participate in the most redundancy-dominated n-plets at all orders, suggesting that they might be the basis of such core.

Interestingly, the core of groups (I1, I2, I3) seems to be broader than the one of the older population, including also modules 9, 10 and 16.

A similar analysis performed over the n-plets with smaller values of O-information shows that the lowest values are attained at orders 6-8, suggesting that this is the characteristic order of synergistic relationships exhibited by the data (see Fig. 3 supplementary materials). Brain areas that consistently participate in synergy-dominated arrangements for all interaction orders include modules 9 and 16. Interestingly, modules that belong to the redundant core tend not to be involved in very synergistic arrangements, with the exception of modules 9 and 16 – both of which abandon the redundant core in the older population.

## 4. Discussion

The present study assessed the high-order redundant and synergistic interactions among brain regions of participants of various ages. Overall, an important increase in redundant interdependencies in the older population was found at all interaction orders. Additionally, a redundant core of brain modules was observed, which decreased in size with age. Together, these two findings suggest a change in the balance of differentiation and integration towards more synchronized arrangements.

We also assessed redundant cores to determine what brain areas were participating in redundancy-dominated arrangements at all interaction orders. The common part which existed in all age groups was composed of modules 2, 5 and 13. While a complete anatomical description of these modules is available in Ref. [29, 33], it is important to emphasize that these three modules have in common the middle frontal gyrus (MFG) and precentral gyrus (PG), two important structures that have a very differentiated function. More specifically, while MFG is part of the prefrontal cortex which mediates executive control and working memory, PG is part of the primary motor cortex, and the two structures are well-known to be affected by aging [3, 34, 35, 36].

Although the redundant role found in this study for the interaction between MFG and PG was obtained from participants being at rest, potentially it reflects the fact that older participants typically compensate motor behavioral deficits by recruiting additional activation of the prefrontal cortex in synchronism with their associated motor areas, while younger subjects only activated motor areas to perform the same tasks [37, 38, 39]. Similarly, it has been also shown that older participants, but not young, recruit the prefrontal cortex while performing purely movement tasks such as the inhibitory motor control, thus relying on more cognitive support for the performance of a motor task that younger participants produce in a more automatic manner, i.e., cognitive penetration into action [40].

The redundant core existing in the young participants – but not in the older participants – was composed of modules 9,10, and 16. The brain structures supporting these modules are composed by the middle and superior frontal gyri, posterior cingulate cortex and the precuneus, all part of the default mode network (DMN), which is an important network of the human brain (see [41, 42] and references therein). Despite other studies have found a lesser participation of the precuneus into the DMN [43], our data confirmed its participation in agreement with other studies [41, 44]. When comparing these results to the ones in Ref. [20], in which a redundant role of the DMN was shown for *n* = 3 across lifespan, we might hypothesize that at high orders of interaction it might break down in older participants, indicating network re-adaptation or anticipation to damage, as it occurs at the onset of other pathologies such as in the early stage of Alzheimer’s disease [45], after concussion [46] or following a multiorgan failure [47].

When looking at modules that participate in synergistic arrangements, we found that modules 9 and 16 are present in all groups of ages. The two modules have in common the posterior cingulate and precuneus that, as explained above, are part of the DMN. Based on these results, we might hypothesize a dual redundant-synergistic role of the DMN, that in older participants, it seems that the redundant contribution gets impaired. But this statement needs further research.

Our analysis goes beyond traditional brain-network approaches that focus on pairwise interactions, considering high-order interactions that can assess redundant and synergistic effects. In doing so, we follow the seminal ideas introduced by Tononi, Sporns, and Edelman [22], which posit that high brain functions might depend on the *co-existence of integration and segregation*. Indeed, while the latter enables brain areas to perform specialized tasks independently of each other, the former serves to bind together brain areas towards an integrated whole for the purpose of goal-directed task performance. A key insight put forward in Ref. [22] is that segregation and integration can coexist, and that this coexistence is measurable by assessing the high-order interactions of neural elements via approaches such as the one used in this study. In the context of aging, it has been shown that the balanced segregation-integration might break down, as the inter-network connectivity increases in older people (reducing segregation), but this increment does not correspond to a better performance, therefore indicating that the reduced segregation is related to neuronal disfunction, possibly due to the reduced inhibitory function found in older adults [48].

The current study has some limitations. First, the accurate quantification of redundancy in large-scale brain networks remains still an open problem [49], but by using the measures of overall correlation strength () and synergy-redundancy balance (Ω) we proposed different metrics that can work out as proxy estimators for synergy and redundancy. Second, different brain parcellations can be used to explore high-order functional interactions in the brain, however, a finer spatial resolution compromises the calculation of all the n-plets to assess the O-Information as the combinatorics becomes huge. Finally, the fact that the values themselves of the O-information suggest a predominance of redundancy might be heavily influenced by the nature of fMRI data, and other results might be seen from other kinds of measurements (e.g. EEG or MEG). What is relevant is the increase of O-information, which is clear evidence pointing towards a change of differentiation-integration balance towards more redundant arrangements.

**Figure 5:**
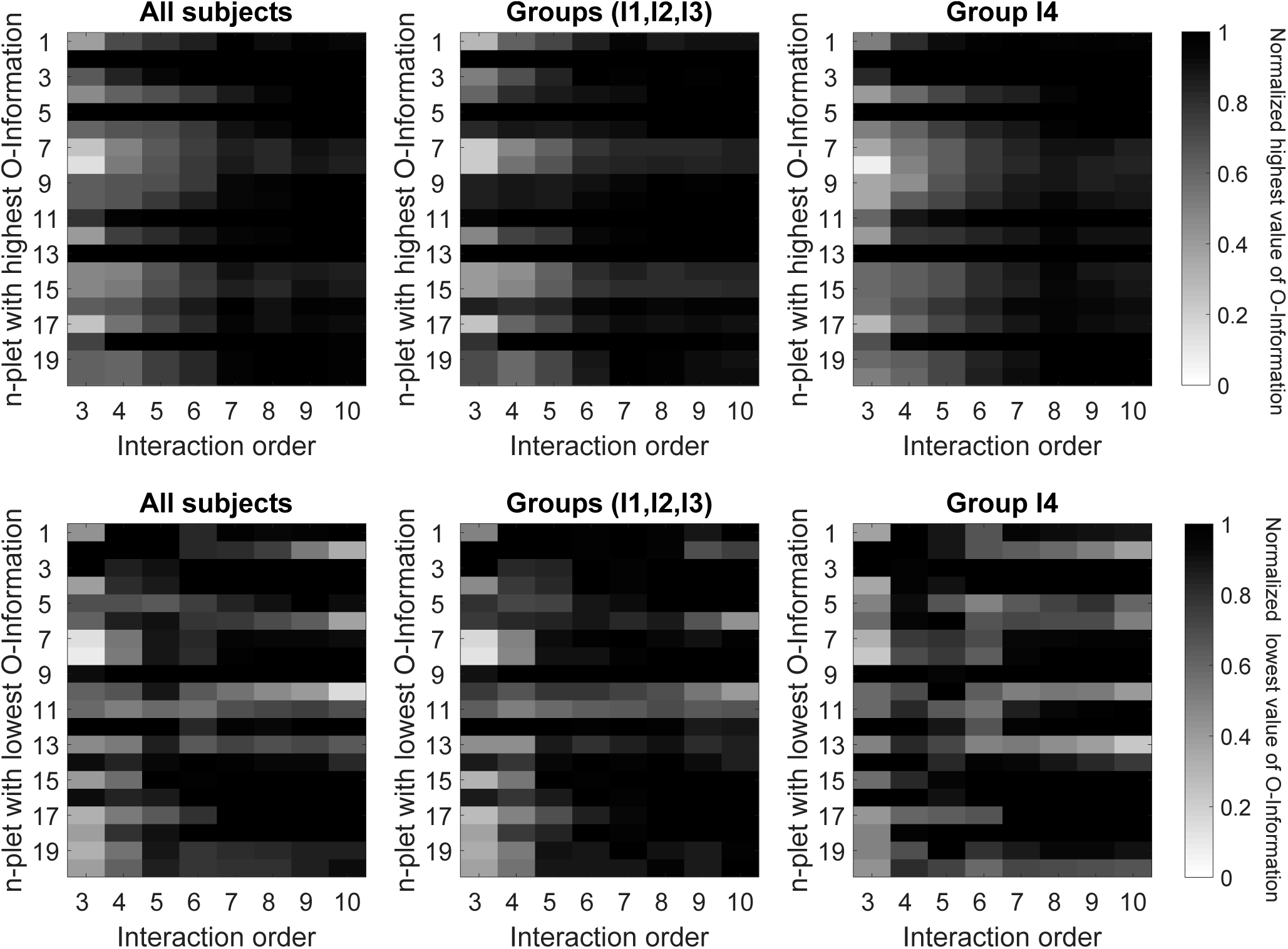
Identification of redundant and synergistic cores along different interaction orders. Top row: As a function of the interaction order *n*, we rank the averaged values of O-Information of all the n-plets per group of participants as detailed in the appendix and illustrated in Fig. 6. We plot for each interaction order the highest value of O-Information in which each module participate, normalized to the n-plet with the highest O-Information averaged in different groups of subjects. For each interaction order the n-plet with the highest values of O-Information appear in black. The highest values of O-Information measure redundancy. Bottom row: The same as before but plotting the lowest values of O-Information, that because all of them were negative, they corresponded to synergy. The two rows show similar plots for three different situations: All subjects (left), when pooling together the three groups (I1, I2, I3) (middle) and for older group I4 (right).

In summary, the framework presented here provide novel insights into the aging brain by assessing high-order functional interdependencies among brain regions, revealing the role of redundancy in prefrontal and motor cortices in older participants, thus affecting basic functions such as working memory, executive and motor functions. We believe that this methodology may help to provide a better understanding of some brain disorders from an informational perspective, providing distinctive patterns of high-order functional behavior or “info-markers”, that may lead to fundamental insights into human brain in health and disease.

The code to compute the metrics used in this article are available at https://github.com/brincolab/High-Order-interactions.

## Acknowledgements

The authors thank Karine Bertin for their valuable comments about our work. M.G. was partially supported by CONICYT-PFCHA/ Doctorado Nacional/ 2019-21190577. R.C. acknowledges finantial support from CON-ICYT PAI Inserción 79160120, Proyecto REDES ETAPA INICIAL, Convocatoria 2017 REDI 170457, Fondecyt Iniciación 2018 Proyecto 11181072. P.M. was funded by the Wellcome Trust (grant no. 210920/Z/18/Z). F.R. acknowledges the support of the Ad Astra Chandaria Foundation. S.P.S was supported by the the FWO Research Foundation Flanders (G089818N), the Excellence of Science funding competition (EOS; 30446199) and the KU Leuven Special Research Fund (grant C16/15/070). J.M.C acknowledges financial support from Ikerbasque (The Basque Foundation for Science) and from the Ministerio Economia, Industria y Competitividad (Spain) and FEDER (grant DPI2016-79874-R), and from the Department of Economical Development and Infrastructure of the Basque Country (Elkartek Program, KK-2018/00032). The Centro Interdisciplinario de Neurociencia de Valparaíso (CINV) is a Millennium Institute supported by Grant P09-022-F of the Millennium Scientific Initiative (ICM-MINECON, Chile)

## Appendix

Here we provide details on how to compute the information-theoretic metrics employed in this work.

**Definition:** The Ω-information of a group of *n* random variables ***X***^*n*^ = (*X*_1_, …, *X*_*n*_) is defined as:

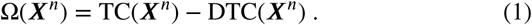

**Definition:** The S-Information (also denoted Σ-information) of a group of *n* random variables ***X***^*n*^ = (*X*_1_, …, *X*_*n*_) is defined as:

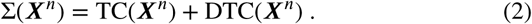

Above, TC and DTC represent the **total correlation** [50] the **dual total correlation** [51] respectively, which are en by:

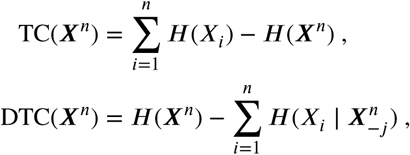

where *H* represent the Shannon entropy and 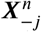 represent the vector of *n* − 1 variables composed by all minus *X*_*j*_ i.e., (*X*_1_,…, *X*_*j*−1_, *X*_*j*+1_,…, *X*_*n*_) Both TC and DTC are non-negative generalisations of mutual information, in the sense that they are zero if and only if all variables *X*_1_, …, *X*_*n*_ are statistically independent of one another. These quantities are computed here using Gaussian Copulas [19].

More formally, we say that [21]:

**Definition:** A system of *n* random variables is redundancy-dominated if Ω(*X*_1_, …, *X*_*n*_) > 0 and synergy-dominated if Ω(*X*_1_, …, *X*_*n*_) < 0.

To compute from the data the O-Information and the S-Information we compute for each subject and for each interaction order the following quantities:

**Figure 6:**
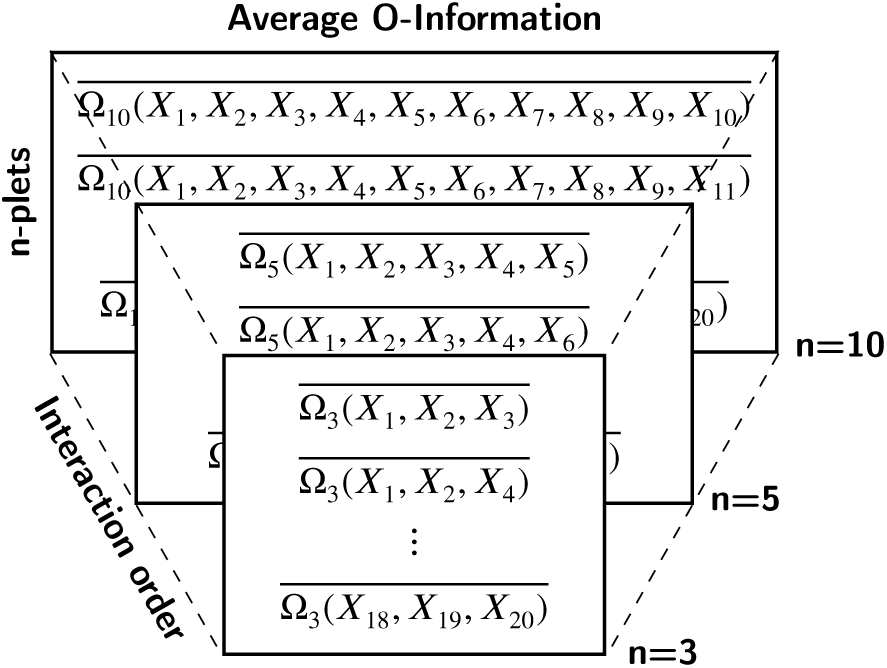
Average O-Information per n-plets over subjects belonging to a certain group. For each subject we compute the O-Information of each n-plet. Then we average the value of O-Information of each n-plet across subjects. These values are the ones depicted in Fig. 5.

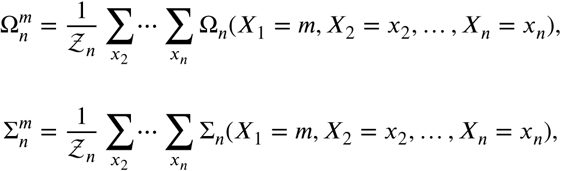

where *m* represents module, *n* interaction order and

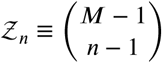

is the total number of subsets of size *n* − 1 in a set of M modules (here, we took M=20). We computed this quantity for all the n-plets {*X*_1_, *X*_2_, …, *X*_*n*_} considering {*x*_1_ = *m* ≠ *x*_2_ ≠ … ≠ *x*_*n*_} with *m* = {1, … *M*}.

These quantities represent the average O-Information and S-Information per subject and brain module. Finally, we average over modules

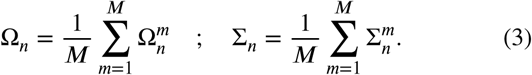

When splitting the values of O-Information on positive and negative values, represented respectively by 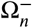 and 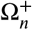, the redundancy and synergy per module and interaction order was computed as:

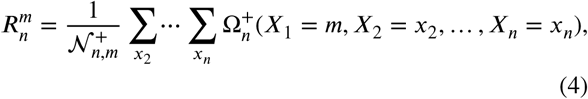

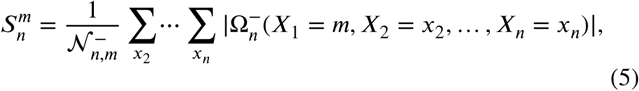

where 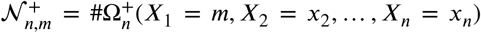 and 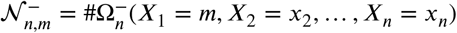 represent the number of positive and negative n-plets for all the n-plets {*X*_1_, *X*_2_, …, *X*_*n*_}, and {*x*_1_ = *m* ≠ *x*_2_ ≠ … ≠ *x*_*n*_} with *m* = {1, …, *M*}. Moreover, the average over all modules was computed as follows:

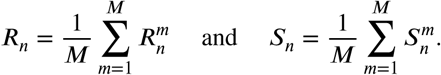

### CRediT authorship contribution statement

**Marilyn Gatica:** Conceptualization of this study, Methodology, Performed the analyses, Made the figures, Drafted the Manuscript. **Rodrigo Cofré:** Conceptualization of this study, Methodology, Performed the analyses, Supervised Information Metrics, Drafted the Manuscript. **Pedro A.M. Mediano:** Conceptualization of this study, Methodology, Supervised Information Metrics, Drafted the Manuscript. **Fernando E. Rosas:** Conceptualization of this study, Methodology, Supervised Information Metrics, Drafted the Manuscript. **Patricio Orio:** Conceptualization of this study, Methodology. **Ibai Diez:** Preprocessed the images. **S.P. Swinnen:** Recruited the participants, Drafted the Manuscript. **Jesus M. Cortes:** Conceptualization of this study, Data curation, Methodology, Supervised Information Metrics, Drafted the Manuscript. All the authors wrote the article and agreed on its submission.

---

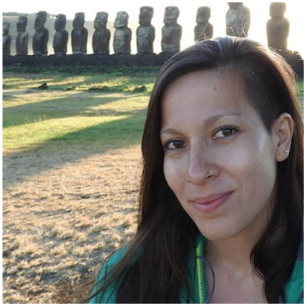

Marilyn Gatica received her Bachelor’s degree in 2013 and her professional degree in Mathematical Engineering in 2014, both from Universidad de Santiago de Chile. She is currently a Ph.D. candidate at Universidad de Valparaíso, Chile and University of the Basque Country (UPV/EHU), Spain. Her main research interests include Computational Neuroscience, Information Theory, Neural Network Models and Biomedicine.

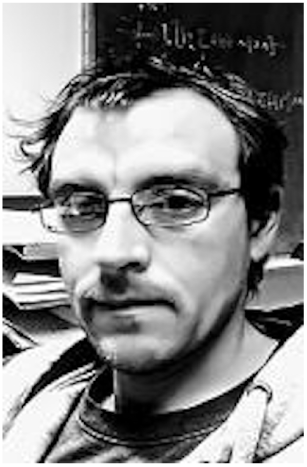

Rodrigo Cofré received his B.Phil in Aesthetics and B.sc. and professional degree in Mathematical Engineering in 2009 from Pontificia Universidad Católica de Chile. He obtained a Ph.D. in Science at the University of Nice (supervised by Prof. Bruno Cessac) and held a postdoctoral position at UNIGE (supervised by Prof. J.-P Eckmann). He is currently joint Professor at the Institute of Mathematical Engineering at Universidad de Valparaíso. His main research interests include complex systems, computational neuroscience and altered states of consciousness.

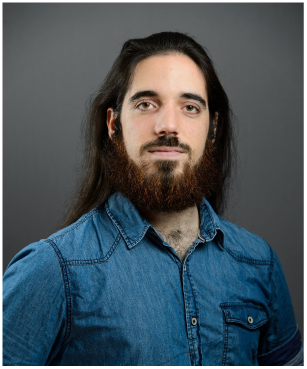

Pedro A.M. Mediano received his Bachelor’s degree in Physics from the University of Valencia, Spain, in 2014, and his PhD from Imperial College London, UK, in 2020. He is currently a postdoctoral researcher in the Consciousness and Cognition laboratory at the University of Cambridge, working on information theory, complex systems, and consciousness neuroscience.

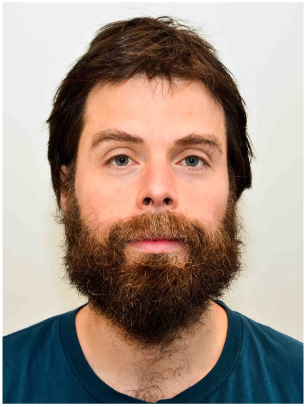

Fernando Rosas received the B.A. degree in music composition and philosophy (Minor), the B.Sc. degree in mathematics, and the M.S. and Ph.D. degree in engineering sciences from the Pontificia Universidad Católica de Chile. He previously worked as Postdoctoral Research Assistant with the Departement Elektrotechniek at KU Leuven, as Postdoctoral Fellow wih the Graduate Institute of Communication Engineering at the National Taiwan University, and as Marie Curie Reseaerch Fellow with the Department of Mathematics and the Department of Electrical and Electronic Engineering at Imperial College London. His research interests lay in the interface between computational neuroscience and complexity science.

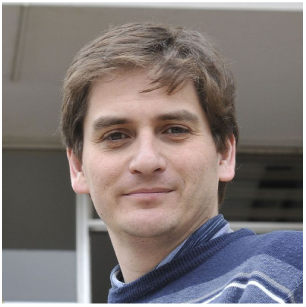

Patricio Orio is a Biochemist and Ph.D. in Molecular and Cell Biology from the Universidad de Chile. He is currently a Full Professor at the Instituto de Neurociencias in the Universidad de Valparaíso and Adjunct Researcher in the Centro Interdisciplinario de Neurociencia de Valparaíso CINV. After an initial training in experimental ion channel biophysics (PhD thesis) and cellular neurophysiology (postdoctoral training), he consolidated an independent career in the field of numerical simulation and analysis of biophysically-inspired models of neural dynamics.

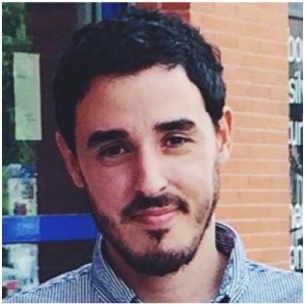

Ibai Diez received his Bachelor’s degree in Telecommunication Engineering in 2009 from Deusto University, Spain, and his PhD in 2015 from University of the Basque Country (UPV/EHU), Spain. He is currently a postdoctoral researcher in Neurology Department at Massachusetts General Hospital – Harvard Medical School. His research interest include structural and functional brain connectivity, biomedical data analysis and functional integration in the brain.

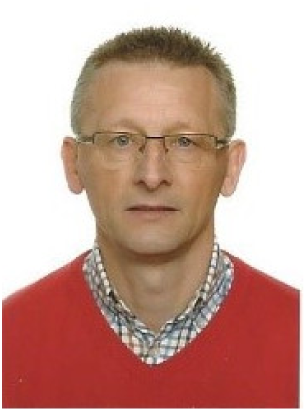

Stephan P. Swinnen started his research career in movement control at the University of California at Los Angeles (UCLA) (1983-85) under the direction of Prof. R. A. Schmidt. He completed his PhD at KU Leuven in 1987. He was awarded a Francqui Research Professor position (2013-2016) and currently directs the Movement Control & Neuroplasticity Research Group at KU Leuven (Belgium). His current research interest is focused on mechanisms underlying movement control and neuroplasticity in normal and pathological conditions using a multidisciplinary approach focusing on the study of brain function, structure, connectivity and neurochemicals.

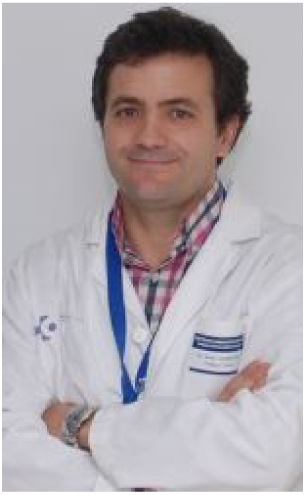

Jesus M. Cortes is an Ikerbasque Research Professor and the head of the Computational Neuroimaging Laboratory of the Biocruces-Bizkaia Health Research Institute in Bilbao (Spain). He teaches Brain Connectivity and Neuroimaging in the M.Sc of Biomedical Engineering. He obtained a Ph.D. in Physics in 2005 and performed three postdoctoral positions in The Netherlands (supervised by Prof. Bert Kappen), USA (supervised by Prof. Terry Sejnowski) and UK (supervised by Prof. Mark van Rossum). His area of research is now focused on brain connectivity, neuroimaging and machine learning methods applied to healthy and pathological conditions.

